# Mice hypomorphic for *Pitx3* define a minimal dopamine neuron population sufficient for entraining behavior and metabolism to scheduled feeding

**DOI:** 10.1101/2020.10.23.353193

**Authors:** Lori L. Scarpa, Brad Wanken, Marten Smidt, Ralph E. Mistlberger, Andrew D. Steele

## Abstract

*Pitx3^ak^* mice lack a functioning retina and nearly all dopamine neurons of the substantia nigra (SN). del Rio-Marten et al (2019) reported that entrainment of circadian rhythms to daily light-dark and feeding schedules is absent in these mice. With food limited to 12h/day, food anticipatory circadian rhythms failed to emerge, and metabolic rhythms failed to synchronize with locomotor and feeding rhythms. The authors propose that retinal innervation of the suprachiasmatic nucleus clock is required for development of cyclic metabolic homeostasis, but methodological issues limit interpretation of the results. Using standardized feeding schedules and procedures for distinguishing free-running from entrained circadian rhythms, we confirm that behavioral and metabolic rhythms in *Pitx3^ak^* mice do not entrain to LD cycles, but we find no desynchrony between these rhythms nor a deficit in entrainment to daily feeding schedules. SN dopamine neurons surviving in *Pitx3^ak^* mice may define a mininum population sufficient for food entrainment.

## INTRODUCTION

Most biological processes display circadian rhythms generated by cell autonomous circadian clocks in the brain and throughout the body (Hastings et al, 2003). A critical function of circadian clocks is to coordinate behavior and physiology with the solar day in accordance with species-specific adaptations to nocturnal or diurnal lifestyles. Synchrony of clocks and rhythms to local environmental time is achieved by adjustments of clock phase in response to periodic stimuli, known collectively as “zeitgebers” (Pittendrigh, 1993). Light is the primary zeitgeber for most species, but non-photic stimuli are also important (Mistlberger and Antle, 2011; Tahara and Shibata, 2018). In mammals, light controls circadian timing via melanopsin-expressing, intrinsically photoreceptive retinal ganglion cells (ipRGC) that project directly to the suprachiasmatic nucleus (SCN) (Berson et al, 2002; Hattar et al, 2002; Güler et al., 2008), a brain structure that serves as a central clock controlling the timing of appetitive behaviors (e.g., sleep-wake, feeding, drinking) and most physiological functions (Golombek and Rosenstein, 2010; Patton and Hastings, 2018). Some SCN-controlled rhythms serve as ‘nonphotic’ time cues that participate in phase control of circadian clock cells elsewhere in the brain and body, and are crucial for coordinating clocks in different tissues with each other and with the solar day (Dibner et al, 2010; Schibler, 2009). Non-photic stimuli of special significance include nutrients and neural and hormonal signals associated with the daily rhythm of food intake.

The importance of feeding for circadian organization in mammals is readily demonstrated by restricting food availability to a limited time of day or night. This shifts the phase of circadian clocks in most peripheral organs and tissues, and induces a daily rhythm of food anticipatory activity superimposed on the LD-entrained rest-activity cycle (Damiola, et al., 2000; Mistlberger, 2011; Patton et al, 2014; Pezuk et al., 2010; Stokkan, et al., 2001; Stephan, 2002). Remarkably, if food is limited to the middle of the light-period in nocturnal mice or rats, peripheral clocks and associated food anticipatory rhythms of activity and physiology also shift, while the light-entrained SCN pacemaker and clocks in a few other tissues (e.g., the pineal gland) do not. Ablation of the SCN eliminates circadian rhythms when food is available ad libitum but does not prevent entrainment of food anticipatory activity and physiology when food is made available on a circadian schedule (Boulos and Terman, 1980; Mistlberger, 1994; Stephan, 2002). This distributed organization of the circadian clock system, combined with tissue-specific sensitivity to different zeitgebers, confers flexibility on the daily circadian program, enabling mammals to reconfigure the system in response to environmental challenges.

The photic input pathway for circadian entrainment has been well-defined, from retinal phototransduction to the signal transduction pathways in SCN neurons that acutely alter the expression of cycling clock genes in response to retinal input, thereby resetting the SCN pacemaker (Dibner, et al., 2010; Golombek and Rosenstein, 2010). Although a number of feeding-related stimuli have been identified that can shift circadian clocks in one or more tissues, the site of the clocks that control food anticipatory behavioral rhythms, and the stimuli that entrain these clocks to daily mealtimes, remain uncertain (Mistlberger, 2011, 2020). Several observations suggest a role for dopamine-sensitive circadian oscillators in the dorsal striatum (Gallardo, et al., 2014a; Hood et al, 2010; LeSauter et al., 2018; Liu, et al., 2012; Smit, et al., 2013). Dopamine-deficient mice lack appetitive behaviors by which to assess food anticipation, but restoration of dopamine signaling in the dorsal striatum is sufficient to rescue food anticipatory activity (Gallardo, et al., 2014). A necessary role for circadian clocks in the dorsal striatum remains to be established.

del Rio-Marten and colleagues (2019) recently reported in this journal that mice harboring a homozygous hypomorphic mutation in the promoter region of *paired-like homeodomain 3 (Pitx3)* exhibit a marked impairment in both LD and food entrainment. *Pitx3^ak^* mutant mice were found to free-run in the presence of LD when food was available ad-libitum. When food was restricted to the 12h dark period, *Pitx3^ak^* mutant mice failed to exhibit food anticipation, and daily rhythms of energy expenditure and locomotor activity oscillated out of phase, and were not synchronized to feeding time. Circadian oscillations of plasma corticosterone and clock gene expression in the ventromedial hypothalamus remained out of phase across mice, although oscillations of clock genes in the liver and brown adipose tissue did synchronize to feeding.

*Pitx3* plays a key role in coordination of gene expression and determination of cell fate in development (Lamonerie, et al., 1996), and is responsible for activating expression of the *tyrosine hydroxylase* gene, which is required for synthesis of dopamine (Cazorla et al., 2000; Lebel, et al., 2001). Expression of Pitx3 is mostly restricted to the eye and midbrain dopamine neurons, with mRNA present 11 days post-conception in the lens (Semina, et al., 1997) and 0.5 days later in the brain, correlating with appearance of midbrain dopaminergic neurons (Smidt et al., 1997). *Pitx3^ak^* mutant mice lack a lens, fail to develop a functional retina and retinohypothalamic tract, and are blind (Rieger, et al., 2001; del Rio-Marten, et al., 2019). In the brain, expression of *Pitx3* is essential for the development of dopamine neurons in the midbrain substantia nigra (SN) but not the ventral tegmental area (VTA) (Smidt, et al., 1997; Nunes, et al., 2003). *Pitx3^ak^* mice exhibit a >90% reduction in dopamine innervation specific to the dorsal striatum (Smidt, et al., 2004; Hwang, et al., 2003). The findings of del Rio-Martin et al that *Pitx3^ak^* mice do not entrain to either LD or feeding schedules are therefore consistent with the known role of the retina in circadian photoentrainment, and with the proposed role for dorsal striatal dopamine innervation in food entrainment. The authors concluded that lack of light input disrupts SCN development and thereby irreversibly impairs cyclic metabolic homeostasis.

While the retinal defects and absence of photic entrainment in *Pitx3^ak^* mice were convincingly demonstrated by del Rio-Marten and colleagues (2019), the methods used to assess entrainment by daily feeding schedules, and to analyze group data in blind mice free-running in LD, raise interpretative issues, which we elaborate below. Here, using the same *Pitx3^ak^* mutant mouse strain, we confirm the absence of photic entrainment and the marked reduction in the number of SN dopamine neurons, but do not observe impaired entrainment of food anticipatory rhythms or metabolism to daily feeding schedules, and do not observe desynchrony between locomotor and metabolic rhythms. The small number of SN dopamine neurons evident in *Pitx3^ak^* mice may define a minimum subset of dopamine neurons sufficient for behavioral and metabolic entrainment to scheduled feeding.

## Results

### Circadian activity and metabolic rhythms free run in *Pitx3^ak^* mice fed ad-libitum in LD

To phenotype behavioral and metabolic circadian rhythms in *Pitx3^ak^* mice, we conducted two experiments using separate cohorts of mutant and control mice. In Experiment 1, young adult *Pitx3^ak^* and control mice (6 male and 6 female per group) were housed individually in metabolic cages (TSE LabMaster/PhenoMaster) to measure feeding, locomotor activity and O_2_ consumption (VO_2_), CO_2_ production (VCO_2_), respiratory exchange ratio (RER), and energy expenditure (EE) by indirect calorimetry. Food was food available ad-libitum for 3 days prior to 28 days of restricted feeding. Data for males and females are presented separately. Sex differences were evident in body weight (males were heavier), fat mass (males were greater), and energy expenditure (females were greater), in both the *Pitx3^ak^* and control groups. Lean mass, food intake per gram body weight, and total daily activity did not differ by sex (Fig. 1; supplementary Fig. S1).

**Figure 1.**
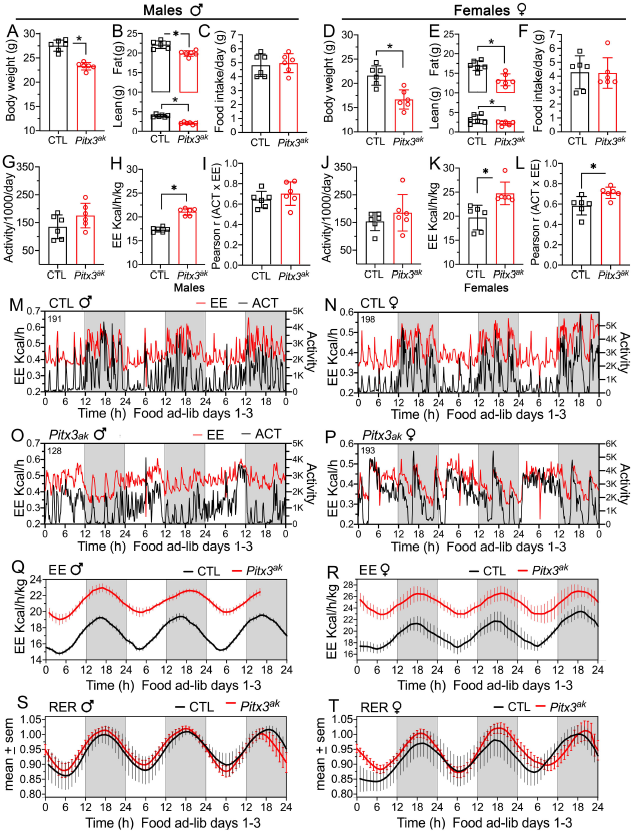
Circadian Rhythms of Activity and Metabolism are Synchronous and Robust in *Pitx3^ak^* Mice Fed Ad-libitum. (A,B,D,E) Body weight and lean and fat mass (grams), in control (CTL, black bars and symbols) and *Pitx3^ak^* groups (red). (C,F) Daily food intake. (G,J) Daily activity counts (divided by 1000). (H,K) Daily energy expenditure (EE, Kcal/h/kg). (I,L) Correlations (Pearson r) between activity (ACT) and EE in 10 min bins across the 3 days of ad-libitum food access for individual mice. (M-P) EE and locomotor activity in 10 min time bins in representative CTL and *Pitx3^ak^* male and female mice, during ad-libitum food access. (Q, R) Group mean hourly EE during ad-lib food access in CTL and *Pitx3^ak^* male and female mice. Data from *Pitx3^ak^* mice were aligned by the first peak and then averaged, and the group waveform was then aligned with the CTL group waveform by the first trough. Data from individual mice were smoothed (4 h running average) before calculating group averages. (S,T) Group mean respiratory exchange ratio (RER). Group means are plotted ± SEM. *p<.05 by ANOVA (Sidak’s post hoc test). Grey shading denotes lights-off.

Compared to control mice, *Pitx3^ak^* mice had lower body mass, fat mass and lean mass (Fig. 1A,B,D,E; Fig. S1). *Pitx3^ak^* mice also ate less, although not when normalized to their body weight (Fig. 1D,F), and showed a non-significant trend for more activity (20.4% and 25.4% higher in male and female *Pitx3^ak^* groups, respectively; p>.05) (Fig. 1G,J). With food available ad-libitum, control mice of both sexes were active and ate predominantly at night, and exhibited daily rhythms of RER, EE, VO_2_, and VCO_2_ with a nocturnal peak (Fig. 1M,N,Q-T; Fig. 2A,B, days AL1-AL3). Individual *Pitx3^ak^* mice also exhibited circadian variations in metabolic variables (e.g., Fig. 1O,P; supplementary Fig. S2), but the rhythms were not synchronized with the LD cycle or with other mice, and thus were not reflected in the group average waveforms (Fig. 2C-F, days AL1-AL3).

**Figure 2.**
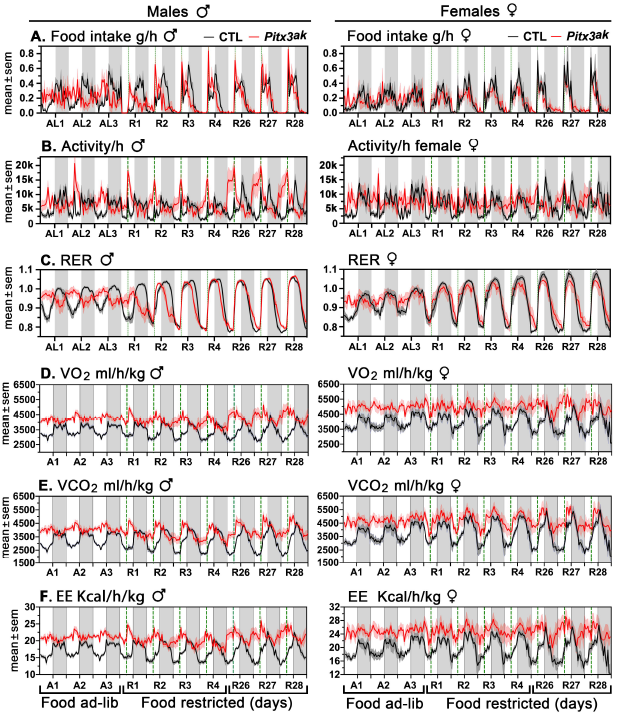
Circadian Rhythms of Activity and Metabolism Synchronize to a Daily Feeding Schedule in *Pitx3^ak^* Mice. Data are group mean (± SEM) hourly averages across the last 3 days of food ad-libitum (A1-3), the first 4 days (R1-4) of restricted feeding (80% of ad-libitum caloric intake provided daily 6h after lights-on), and the last 3 days (R26-28) of restricted feeding. (A) Food intake (grams), (B) locomotor activity (where k denotes thousand), (C) respiratory exchange ratio, (D) oxygen consumption, (E) carbon dioxide production, and (F) energy expenditure. Grey shading denotes lights-off. Control (CTL) groups are coded black and *Pitx3^ak^* groups are coded red. Male groups are in the left column and female groups in the right column.

To compare metabolic rhythms between groups, data from individual *Pitx3^ak^* mice (e.g., Fig. S2) were aligned by the onset of the daily active period, smoothed (4 h running average) and then averaged across mice, separately for males and females. The *Pitx3^ak^* group average waveforms were then aligned with average waveforms from the control mice. Overlays of these waveforms clearly show that circadian variations of RER (Fig. 1E,F) and EE (Fig. 1G,H) in *Pitx3^ak^* mice were very similar to control mice, despite the lack of entrainment to LD. Circadian variations of VO_2_ and VCO_2_ closely paralleled EE, as expected (data not shown). The primary difference between groups was the markedly elevated EE, VO_2_ and VCO_2_ throughout the circadian cycle in the *Pitx3^ak^* mice of both sexes. This is likely due in part to the significantly lower body mass and somewhat elevated activity levels in the *Pitx3^ak^* mice.

Importantly, metabolic activity and locomotor activity were highly correlated in all individual mice, and there was no evidence for a lack of synchrony between locomotion and EE or O_2_ consumption in *Pitx3^ak^* mice, as reported by del Rio-Martin et al (2019). The tight coupling between metabolism and locomotion in *Pitx3^ak^* and control mice is clearly evident in running waveforms of EE and activity counts in 10 min time bins for representative mice in each group (Fig. 1M-P). Correlation coefficients between EE and locomotor activity averaged at least +.60 in each group, were statistically equivalent in the two male groups, and were significantly higher in the *Pitx3^ak^* females compared to the control females (Fig. 1I, L).

### Circadian metabolic rhythms synchronize to daily feeding schedules in *Pitx3^ak^*

In adult C57BL/6J mice maintained in constant dark or dim light, feeding schedules that provide a reduced amount of calories delivered at a fixed time once every 24h typically do not entrain free-running activity rhythms mediated by the SCN pacemaker, but do induce a daily rhythm of food anticipatory activity controlled by food-entrainable circadian oscillators located outside of the SCN (Boulos and Terman, 1980; Mistlberger, 1994; Stephan, 2002). del Rio-Martin et al (2019) reported that *Pitx3^ak^* mice fed only during the 12h dark period failed to exhibit a food anticipatory activity rhythm, and failed to synchronize metabolic rhythms with the 24h feeding schedule or with concurrent circadian cycling of locomotor activity. To evaluate entrainment of anticipatory activity rhythms and metabolism to daily feeding schedules, *Pitx3^ak^* and control mice in Experiment 1 were restricted to 80% of group mean ad-libitum caloric intake, provided each day 6h before lights-off, for 28 days. Scheduling meal onset in the middle of the light period is the conventional method for dissociating food anticipatory activity rhythms from nocturnal activity controlled by the SCN pacemaker. The mice were continuously monitored in the metabolic cages on days 1-4 and 26-28 of food restriction.

Group mean and individual waveforms of hourly (Fig. 2A), 10 min (Fig. 3I-L), and cumulative food intake (Fig. 3A,C) indicate that *Pitx3^ak^* and control mice consumed 85-95% of the daily food allotment within ~12h, with a range of 8h to 22h for individual mice across days, and no group differences. All the mice began eating immediately at food delivery. Most control mice showed a second peak of food intake ~6h later, at lights-off (Fig. 2A). Some *Pitx3^ak^* mice also showed a second peak ~5-6h after food delivery. By day 28 of food restriction, body weight in male mice was reduced by 7.0 ± 1.3 % in the *Pitx3^ak^* group and 4.2 ± 1.6% in the control group (between groups, *t_10_* =4.05, *p*=.0023; Fig. 3B). Weight loss in female mice was more variable and not significant for either the *Pitx3^ak^* (0.2 ± 4.8 %) or the control (1.8 ± 2.2 %) groups (Fig. 3D). Two the control females and three *Pitx3^ak^* females gained weight across the 28-day feeding schedule. This may be a consequence of using group mean ad-libitum caloric intake to determine the amount of food provided for individual mice during restriction.

**Figure 3.**
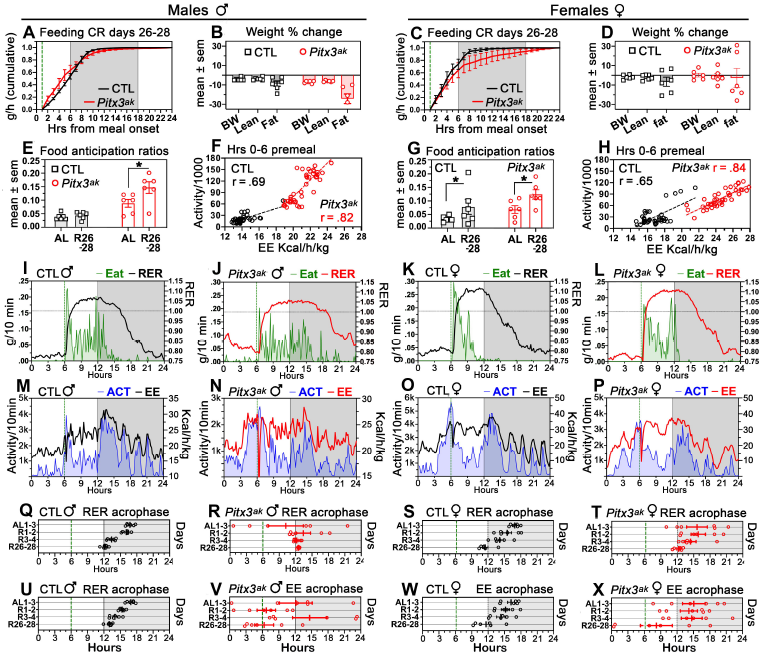
Circadian Rhythms of Activity and Metabolism Synchronize to a Daily Feeding Schedule in *Pitx3^ak^* Mice. (A,C) Group mean (±SEM) cumulative hourly food intake averaged across the last 3 days of restricted feeding (R26-28) in control (black) and *Pitx3^ak^* mice (red). (B,D) Percent change of body weight (BW), lean and fat mass after 28 days of restricted feeding. (E,G) Food anticipation ratios averaged over the last 3 days of restricted feeding (R26-28) and the last 3 days of ad-libitum food access (AL) for statistical comparison. * denotes p<.05 (Sidak’s post hoc test). (F,H) Correlation between activity and energy expenditure during the 6 h prior to mealtime on the last 3 days of restricted feeding (Pearson correlation coefficient r). (I-L) Food intake (green line and shading) and respiratory exchange ratio (RER) per 10 min time bin averaged over the last 3 days of restricted feeding in representative male and female control and *Pitx3^ak^* mice. (M-P) Locomotor activity and energy expenditure (EE) per 10 min time bin averaged over the last 3 days of restricted feeding in the same representative mice. (Q-T) Acrophase of the daily rhythm of RER for individual mice averaged over ad-libitum food access days (AL) 1-3, and restricted feeding days R1-2, R3-4, and R26-28. (U-X) Acrophase of the daily rhythm of RER for individual mice. Grey shading denotes lights-off.

Group mean waveforms of activity during restricted feeding reveal increased activity prior to mealtime on days 26-28 of caloric restriction in the *Pitx3^ak^* male and female groups, and in the control female group (Fig. 2B). Waveforms from some individual mice show activity beginning to rise ~ 3h prior to mealtime and rising to a peak at mealtime (Fig. 3N-P), although this pattern was weak or absent in some other mice (e.g., Fig. 3M). Food anticipatory activity was quantified as a ratio of activity during the 3h prior to mealtime relative to total daily activity. For statistical analysis, anticipation ratios were averaged over the last 3 days of restricted feeding and compared with ratios calculated for the 3 days of ad-libitum food access. Two-way repeated measures ANOVA confirmed a significant main effect of restricted feeding (*F_1,20_* = 32.29, *p* < .0001) and group (*F_3,20_* = 9.859, *p* =.0003), and a significant interaction (*F_3,20_* = 3.226, *p*=.044). Post hoc tests (Sidak’s) comparing food restriction with ad-libitum days indicate significant anticipation in the male and female *Pitx3^ak^* groups, and in the female control group (Fig. 3E,G). Low food anticipation ratios in male control mice and some mice in the other groups likely reflects the prolonged duration of food intake and minimal weight loss in these cases.

Despite low food anticipation in many of the mice, circadian variation of RER synchronized to the daily feeding schedule in all mice. The effect is particularly striking in the average waveforms for the *Pitx3^ak^* male and female groups, which showed no group rhythm when food was available ad-libitum, but rapid emergence of group synchrony when food was restricted (Fig. 2C; Fig. 3I-L). For statistical analysis, RER rhythm timing for each mouse was represented by the acrophase of a cosine function fit to each day of recording, and then averaged across the 3 days of ad-libitum food access, and the last 3 days of restricted feeding. Repeated measures ANOVA confirmed a significant main effect of restricted feeding (*F_1,20_* = 8.804, *p* = .0076) but no main effect of group (*F_3,20_* = 2.157, *p* = .125). Post hoc tests indicated no differences in RER acrophase between control and *Pitx3^ak^* groups, for males or females.

The circadian rhythm of EE in control male and female mice showed a similar phase advance shift in response to the feeding schedule (Fig.2F; Fig.3U,W; Fig. S3). In the *Pitx3^ak^* groups, the EE rhythm acrophase also shifted toward meal onset, but was considerably more variable across mice (Fig. 3V,X). Differences within the *Pitx3^ak^* groups, and between the *Pitx3^ak^* and control groups, appear to reflect differences in the amount of activity prior to mealtime. Visual inspection of the group mean waveforms for locomotor activity and EE (Fig. 2B, F) clearly shows the high degree of synchrony between metabolic rhythms and behavior in all groups. This is further illustrated in plots of activity and EE from representative individual mice (Fig. 3M-P), and in bivariate plots of activity and EE during the 6h prior to mealtime (Fig. 3F,H). The rhythms of VO_2_ and VCO_2_ (Fig. 2D) were highly correlated with both EE and activity. There was no evidence for a lack of synchrony between locomotor activity and metabolic rhythms in *Pitx3^ak^* mice, whether food was available ad-libitum or restricted to 12h/day.

### Circadian activity rhythms are robust and free run in *Pitx3^ak^* mice fed ad-libitum in LD

Activity and metabolic rhythms in *Pitx3^ak^* mice in Experiment 1 did not entrain to the daily LD cycle. Also, during restricted feeding, some mice did not lose weight, and thus showed little food anticipatory activity. Experiment 2 had two objectives. The first was to record mice continuously for many weeks, to measure the period (τ) and phase of free-running circadian activity rhythms in *Pitx3^ak^* mice, and thereby more clearly distinguish food anticipatory activity from free-running locomotor activity. The second objective was to adjust the level of caloric restriction to promote expression of food-entrained anticipatory activity.

Young, female, age-matched *Pitx3^ak^* mice (N=7) and control mice (N=8) were housed individually in cages equipped with photobeams. With food available ad-libitum, locomotor activity was 78 ± 6 % nocturnal in control mice (Fig. 4A), but free-ran with a periodicity in the 23.25 h to 23.83 h range in *Pitx3^ak^* mice (mean ± SD, 23.56 ± 0.24 h; Figs. 4B, 5B-F). To compare activity rhythm parameters between groups, the time series for each *Pitx3^ak^* mouse was folded modulo (⍰) (Fig. 4B), and then averaged across the ad-libitum food access days. Average waveforms for each mouse were then aligned by the onset of the daily active period (⍰) to generate a group mean waveform (Fig. 4C; supplementary Fig. S3). Compared to control mice, *Pitx3^ak^* mice averaged 52% more activity counts per day (*t_13_*=3.26, *p*=.0062; Fig. 4D) and exhibited fewer transitions between rest and activity states (lower intradaily variability; *t_13_* = 3.51, *p*=0039; Fig. 4H). There were no significant group differences in ⍰ duration (Fig. 4E), the ratio of activity in the active period to activity in the rest period (⍰ ⍰ ⍰ ratio; Fig. 4F), or the relative amplitude (the ratio of activity in the most active 10h to activity in the least active 5h; Fig. 4G). Apart from the absence of entrainment to LD, circadian regulation of activity with food available ad-libitum was robust in *Pitx3^ak^* mice.

**Figure 4.**
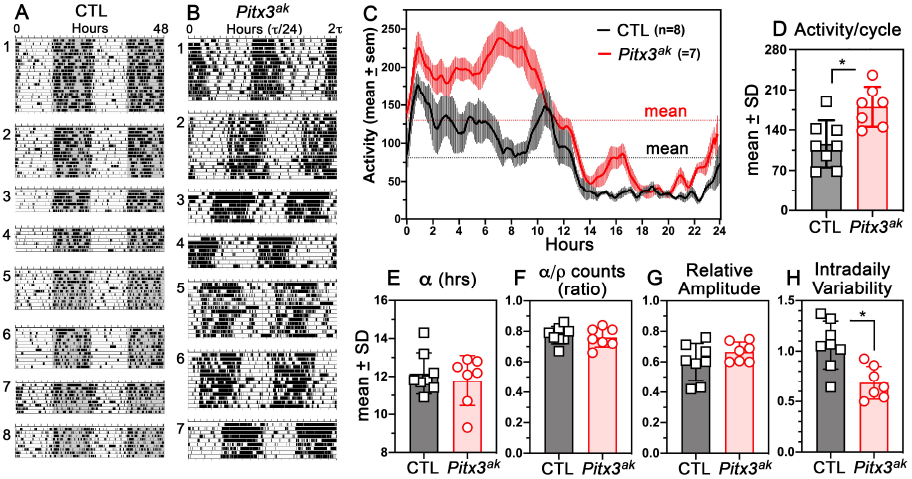
Circadian Activity Rhythms are Robust and Free run in *Pitx3^ak^* Mice Fed Adlibitum in LD. Actograms of locomotor activity prior to restricted feeding in (A) 8 female control mice (CTL), and (B) 7 female *Pitx3^ak^* mice. Activity data are plotted in 10 min times plotted from left to right within a day, and consecutive days are aligned both vertically and horizontally (the graphs are ‘double-plotted, so that each line represents 48h). Grey shading denotes lights-off. The time scale for each *Pitx3^ak^* mouse was set to the period (⍰) of the free-running rhythm, as determined by F-periodogram analysis. (C) Group mean (±SEM) waveform of locomotor activity from the 8 CTL mice (black curve) averaged across 24h days, and from the 7 *Pitx3^ak^* mice (red curve) averaged across circadian cycles (of ⍰ ⍰duration) for each mouse, and then averaged across the group after aligning the individual educed waveforms by the first peak of activity. (D) Group mean (±SD) of activity counts per 24h or circadian cycle. (E) The duration of the daily active period, ⍰. (F) The ratio of activity in ⍰ to activity in the rest phase, ⍰ ⍰ (G) The relative amplitude (RA) of the activity rhythm. (H) The intradaily variability of the rest-activity rhythm. *p<.05 by ANOVA (Sidak’s post hoc test).

### Circadian anticipatory activity rhythms entrain to scheduled feeding in *Pitx3^ak^* mice

To further evaluate entrainment of activity rhythms by scheduled feeding, *Pitx3^ak^* and control mice were provided a restricted amount of food once daily, beginning 6h prior to lights-off, for 35-55 days. In our previous work with adult C57BL/6J mice, a 60% caloric restriction schedule resulted in weight loss in the 10-20% range, and reliably induced a robust circadian rhythm of food anticipatory activity (e.g., Gallardo et al., 2014b). In preliminary work with *Pitx3^ak^* mice, 60% caloric restriction led to excessive weight loss and mortality (supplementary Fig. S4). Caloric restriction adjusted to 80% for the *Pitx3^ak^* mice and maintained at 60% for control mice resulted in equivalent weight loss in the two groups (8.4 ± 3.6 % and 16.9 ± 2.5 %, respectively; p>.05).

Under this feeding regimen, all control mice exhibited robust food anticipatory activity within the first week of restricted feeding (e.g., Fig. 5A1). Activity began to rise ~3h prior to mealtime and increased monotonically to a peak at expected mealtime (e.g., Fig. 5A2, A3). Nocturnal activity decreased but remained synchronized to the LD cycle. Total daily (24h) activity during restricted feeding decreased by 25.7 ± 6.6 % (Fig. 5K; paired *t_7_*=4.797, *p*=.002)

**Figure 5.**
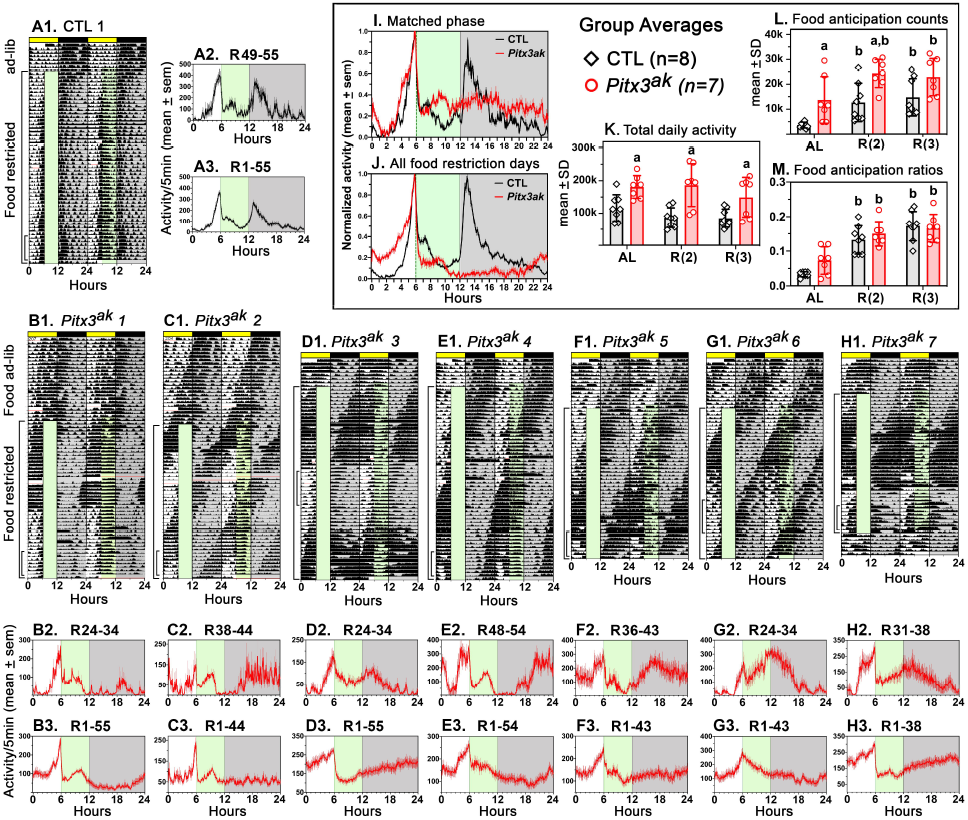
Circadian Anticipatory Activity Rhythms Entrain to Scheduled Feeding in *Pitx3^ak^* Mice. (A1) Double-plotted actogram of locomotor activity in 10 min time bins from a representative control mouse with food available ad-libitum and then restricted to 60% of free-feeding caloric intake, provided 6h after lights-on each day for 55 days. Green shading denotes the first 6h of feeding, but feeding was not time-limited. (A2) Average waveform for days 49-55 of restricted feeding in this mouse. (A3) Average waveform for days 1-55 of restricted feeding for this mouse. (B1-H1) Actograms for all days of recording in the 7 *Pitx3^ak^* mice. (B2-H2) Activity waveforms for 6-10 days when the free-running rhythm did not overlap with the hours before mealtime. (B3-H3) Activity waveforms for all days of food restriction. (I) Group mean waveforms created by normalizing individual waveforms from *Pitx3^ak^* mice in panels B2-H2 (red curve), and in all 8 control mice (A2 and 7 others not shown), and then normalizing the group mean waveforms to emphasis the shape of the waveforms. (J) The same as panel I, but using data from panels A3 and B3-H3. (K) Group mean (±SD) total daily activity in control (black) and *Pitx3^ak^* (red) mice during ad-libitum food access (AL) and during the sets of restricted feeding days represented in panels A2-H2 (labeled R(2)) and A3-H3 (labeled R(3)). (L) Activity during the 3h prior to meal onset. (M) Ratios of activity during the 3h prior to meal onset relative to total daily activity. In the bar graphs, ‘a’ denotes significant (p<.05) differences between groups, and ‘b’ denotes significant differences between restricted feeding days and ad-libitum food access days within groups.

Activity rhythms in *Pitx3^ak^* mice continued to free-run during restricted feeding, but a bout of activity anticipating mealtime was clearly evident in the actograms when mealtime occurred during the rest phase of the free-running rhythm (Fig. 5B1-H1). Activity profiles generated by averaging across a week of these days for each mouse confirmed significant food anticipation beginning 2-3h prior to mealtime and rising to a peak at mealtime (Fig. 5B2-F2). Averaging across all days of restricted feeding completely or substantially removed the contribution of the free-running activity rhythm to the activity profiles, and highlighted mealtime as the determinant of a 24h rhythm of activity in these mice (Fig. 5B3-F3).

The overlay of normalized group mean activity waveforms from the *Pitx3^ak^* and control mice further highlights the similar timing and duration of the food anticipatory rhythm in the two groups (Fig. 5I,J). Compared to control mice, *Pitx3^ak^* mice showed significantly more activity counts during the 3h prior to mealtime (*t_13_* = 3.282, *p*=.006; Fig. 5L), but there was no group difference when anticipatory activity was expressed as a ratio relative to total daily activity (*t_13_* = 0.876, *p*=.3967; Fig. 5M). Unlike control mice, the *Pitx3^ak^* mice did not show a significant reduction in total daily activity during restricted feeding (*t_6_* = =0.31, *p*=.76), and activity levels remained higher than in controls (*t_13_* = 3.89, *p*=.0019; Fig. 5K)

### Visualizing the nigrostriatal dopamine population sufficient for food and metabolic entrainment in Pitx3^ak^ mice

Catecholamine neurons were visualized by immunolabeling for TH (Fig. 6). As expected, in sagittal sections the midbrain and forebrain staining appeared very strong in control mice, showing dense innervation of the striatum, nucleus accumbens, and olfactory tubercle (Fig. 6A). In contrast, there was substantially less innervation of the striatum and nucleus accumbens in the *Pitx3^ak^* sections, whereas the olfactory tubercle appeared strongly stained (Fig. 6B). We stained consecutive coronal sections of the midbrain with TH antibody to visualize the dopamine neurons in the SN and VTA. Control midbrains showed numerous dopamine neurons in the VTA and SN (Fig. 6C) while the *Pitx3^ak^* midbrain showed very little TH staining in the SN but ample staining in the VTA (Fig 6D). Reference atlas pictures from Franklin and Paxinos (2007) are showed in Fig. 6E.

**Figure 6.**
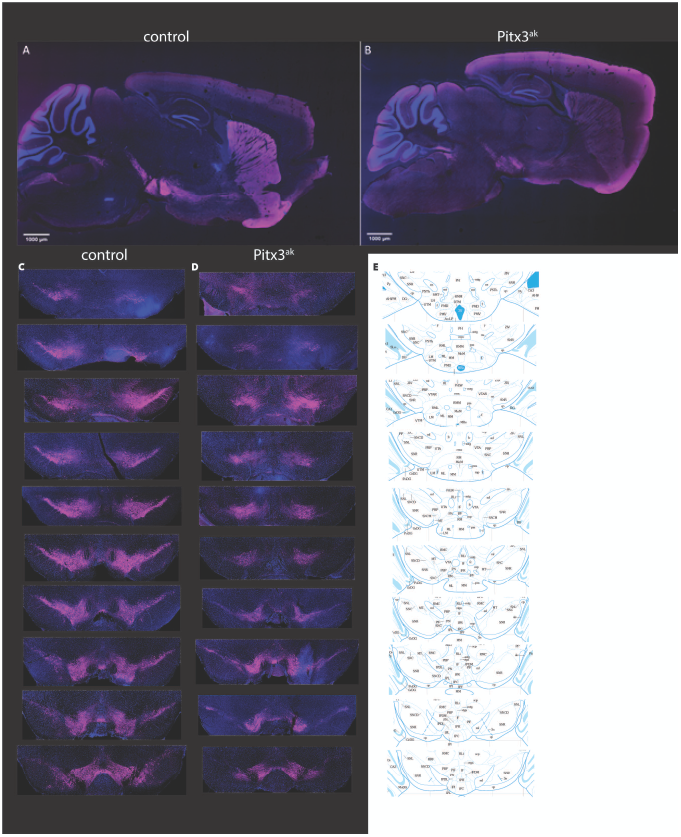
Tyrosine Hydroxylase Antibody Labeling. (A) sagittal section of control brain and (B) *Pitx3^ak^* substantially less innervation of the striatum and nucleus accumbens in Pitx3ak sections (Fig. 6B). We stained consecutive coronal sections of the midbrain with TH antibody to visualize the dopamine neurons in the SN and VTA. Control midbrains showed numerous dopamine neurons in the VTA and SN (Fig. 6C) while the Pitx3ak midbrainshowed very little TH staining in the SN but ample staining in the VTA.

## DISCUSSION

del Rio-Martin et al (2019) reported that mice bearing the *Pitx3^ak^* mutation free-run in the presence of 24h LD cycles, and fail to entrain locomotor activity and metabolic rhythms to a daily feeding schedule. Our results confirm the absence of LD entrainment, but clearly show that the *Pitx3^ak^* mutation does not impair entrainment to feeding schedules, or synchrony between daily rhythms of activity and metabolism.

The absence of food anticipatory activity in the del Rio-Martin study was likely due in part to the extended duration of daily food access (12h/day). Previous studies have shown that food anticipation is weak or absent when daily meal duration is as long as 12h (e.g., Stephan and Becker, 1989; LeSauter et al., 2018), and this may account for weak food anticipation in our Experiment 1, particularly in the control mice. Another factor that may have contributed to weak food anticipation in the del Rio-Martin study was scheduling of food access to the lights-off period. Most studies of food anticipation provide food in the middle of the light period, when activity levels are normally low in control animals, and emergence of anticipatory activity can be clearly distinguished from nocturnal activity. Using this procedure, and restricting calories sufficiently to achieve ~10% or greater weight loss, we were able to demonstrate robust food anticipation in all control and *Pitx3^ak^* mice. Although del Rio-Martin et al attribute an apparent deficit in food anticipation to the *Pitx3^ak^* gene mutation, their control mice also did not show food anticipation.

The feeding schedule used by del Rio Martin may similarly explain the failure of metabolic rhythms to synchronize to daily feeding in their *Pitx3^ak^* mice, but data averaging is also a critical issue. *Pitx3^ak^* mice cannot entrain to LD cycles, and free-run with a circadian period that varies across individuals. It is therefor not illuminating to take averages of data collected from groups of *Pitx3^ak^* mice without controlling for differences in circadian phase at the time of data collection. Averaging data from free-running animals will most likely yield something that approaches a flat line, as we also show here. With food available ad-libitum, we were able to align the free-running activity and metabolic rhythms in groups of *Pitx3^ak^* mice to a common phase (e.g., locomotor activity onset or the peak of the metabolism rhythm), and show that circadian organization is robust, and that activity and metabolism are not dissociated. When food was restricted, rhythms in the metabolic variables quickly emerged in the group data without the need for alignment, indicating that individual *Pitx3^ak^* mice were synchronizing to mealtime, and that there was no deficit in food entrainment, and again no dissociation between activity and metabolic rhythms.

del Rio-Martin et al conclude that failure of the retina to develop and innervate the SCN pacemaker in *Pitx3^ak^* mice permanently impairs cyclic metabolic homeostasis. Our results do not provide support for this model, at least with respect to coordination of behavioral and metabolic rhythms with feeding rhythms. An important role for retinal ipRGCs in development of brain circuits that convey metabolic signals to the SCN pacemaker has recently been suggested based on evidence that selective ablation of ipRGCs or early optic enucleation impairs development of a pathway from the retinorecipient thalamic intergeniculate leaflet to the SCN, and reduces the expression of food anticipatory activity (Fernandez et al, 2020). Given the robust food anticipatory rhythms evident in *Pitx3^ak^* mice, these mice will be a useful model to further evaluate a proposed role for the retina in the development of neural circuits that regulate entrainment of behavior and physiology to daily cycles of food intake.

Several lines of evidence point to an important role for dopamine signaling in the dorsal striatum for induction or expression of food anticipatory circadian activity rhythms, but which and how many of the dopamine neurons are necessary for this rhythm are not known (Liu et al. 2012, Smit et al, 2013, Gallardo, et al., 2014a, Michalik et al., 2015,i LeSauter et al 2018). *Pitx3^ak^* mice develop only a small fraction of the normal complement of substantia nigra dopamine neurons responsible for dopamine signaling in the dorsal striatum, and therefore we were fully expecting to see a significant deficit in food entrainment. Failure to observe any deficit despite a >90% reduction in the number of SN dopamine neurons came as a surprise, and presumably attests to the robust compensatory mechanisms known to be induced in the dorsal striatum in this and other models of chronic dopamine depletion (Hwang et al., 2005; Sagot et al., 2018; Wang et al., 2017). Our results may define a minimum population of SN dopamine neurons sufficient for food entrainment. Further genetic identification of this population is warranted to fully delineate the role of dopamine signaling in food-entrained metabolic and activity rhythms.

## Author Contributions

Conceptualization, L.L.S., R.E.M., A.D.S;

Methodology, L.L.S., B.W., M.S., R.E.M., A.D.S;

Investigation, L.L.S., B.W., M.S., R.E.M., A.D.S;

Writing – Original Draft, L.L.S., R.E.M., A.D.S.;

Writing – Review & Editing, L.L.S., B.W., M.S., R.E.M., A.D.S;

Funding Acquisition, A.D.S;

Resources, B.W., M.S., A.D.S;

Supervision, A.D.S.

## Declaration of Interests

The authors declare no competing interests.

## STAR Methods

### Ethics Statements

All experiments involving the use of animals were approved by the Institutional Animal Care and Use Committee of California State Polytechnic University, Pomona, CA.

### Mouse Strains, Genotyping, and Husbandry

*Pitx3^ak^* mice (strain 000942) were obtained from Jackson labs cryorepository. The aphakia allele arose spontaneously in a 129S1/Sv-p+ Tyr+ KitlSl-J/J strain that was subsequently backcrossed to C57BLKS/J and then to C57BL/6J. Genomic DNA samples were isolated from proteinase K-digested tail tissue using an isopropanol precipitation. Pitx alleles were amplified using the following primers: ‘OR-72F’ CTCTCCAGCCTCCCTCAAATACT, ‘OR-75R’ TCGGATTTGGCTTCTGATGGTTTT, ‘Pitx3-set3-F’

ATTAGAGGTCGTTCAGGATG, and ‘Pitx3-set3-R’ TAATTGAGGCCTTGGGCTCT. PCR cycling conditions were as follows: 95°C for 5 minutes, 35 cycles at 94°C for 30 seconds, 58°C for 30 seconds, and 72 °C for 1.5 minutes followed by 1 cycle at 72°C for 10 minutes. Results were visualized on a 2% agarose gel. *Pitx3-cre; FloxTH* mice were genotyped by Transetyx (Cordova, TN) from tail samples.

All mice were maintained on a 12:12 light:dark cycle throughout all experiments. All mice were single housed in static microisolator cages containing sani-chip bedding (Envigo, 7090) and a cotton nestlet. Room temperature was maintained at 22-24°C with humidity at 20-45%.

### Food Intake and Caloric Restriction Conditions

Mice in Experiment 1 were single housed at 9-10 weeks of age and subjected to a 3-day acclimation period prior to measuring food intake. Food was available ad libitum and daily consumption was calculated from differential weights over 24-, 48- and 72-hours. For caloric restriction (CR) studies, mice were fed 80% of their average daily food intake for a minimum of 28 days, with food delivered in the middle of the light period (6h after light onset).

Throughout the course of the study, weekly measurements of body weight were done on mice receiving 80% CR while mice on 60% CR were measured three times per week.

### Metabolic measurements

In Experiment 1, control and *Pitx3^ak^* mice (n=6 males and females per group) were studied at Children’s Hospital Los Angeles/USC for body composition, metabolic, and behavioral entrainment to scheduled feeding. The EchoMRI-700 Whole Body Magnetic Resonance Analyzer System was utilized for measurements of body fat, lean mass, body fluids and total body water. The TSE LabMaster/PhenoMaster Metabolic Phenotyping System was used to collect data on energy expenditure by indirect calorimetry, feeding behavior and physical activity. Parameters were measured for 7 consecutive days (3 days on ad libitum, 4 days on CR), then again for 3 additional days on days 26 to 28 of 80% CR. Circadian timing of metabolic variables measured in Experiment 1 was quantified by calculating the acrophase of a sine wave fit to each day of recording, using Clocklab (Actimetrics, IL).

### Measurement of home-cage activity

In Experiment 2, a modified Comprehensive Lab Animal Monitoring System from Columbus Instruments (Columbus, OH.) was used to monitor activity continuously with measurements collected in bins of 5 minutes based on the number of infra-red beam breaks across x-,y-, and z-axes. Activity was visualized using ClockLab software (Actimetrics, Il) and GraphPad Prism 8 (GraphPad Software, San Diego CA). Circadian periodicity in the activity data from Experiment 2 was quantified using the F-periodogram as implemented in Clocklab.

### Tissue Histology and Antibody Labeling

Transcardiac perfusion was performed by injecting 5-10 mL of phosphate buffer into the left ventricle followed by 5 mL of 4% PFA (Sigma), both made fresh prior to perfusion. Whole brain tissue was removed and stored in 4% PFA at -4C for 24 hours, then placed in 0.1M PBS. 50-micron coronal and sagittal sections were obtained using a Leica VT1000S Vibratome (Leica Biosystems, Buffalo Grove, IL) and stored in 0.1M PBS at -4°C.

Tissue samples were placed in glass well plates in 0.1M PBS for 1 minute, then permeabilized by incubating in PBST (0.5% Triton X-100 in PBS) for 10 minutes. Samples were then blocked in 2% BSA in goat serum for 10 minutes. Tissue samples were diluted in a tyrosine hydroxylase primary antibody (Aves Lab, Tigard, OR; Catalog #TYH) diluted in goat serum overnight then washed three times in PBS for 5 minutes each. Tissue was submerged in a 1:500 dilution of secondary antibody (ThermoFisher, Catalog #A-21449) to PBS for 1 hour. The tissue samples were washed in PBS two times for 5 minutes each then added to a DAPI solution for 5 minutes. Tissue samples were washed again for 5 minutes in PBS before being mounted.

Imaging was performed using a Nikon C2 Confocal scope with a standard detector system and Nikon Elements Software. Adjustments to images were performed using Fiji (formerly known as ImageJ).

### Statistical Testing

Statistical significance of group differences was evaluated by ANOVA or t-tests conducted using GraphPad Prism 8. Graphs were prepared using GraphPad Prism and figure layouts created using Adobe Illustrator.

## Notes

### Competing Interest Statement

The authors have declared no competing interest.

