## Supplemental Figures for "Mice hypomorphic for *Pitx3* define a minimal dopamine neuron population sufficient for entraining behavior and metabolism to scheduled feeding"

**Fig S1. Sex differences in body mass and composition in control (CTL) and *Pitx3^ak^* mice with food available ad-libitum (Experiment 1).**

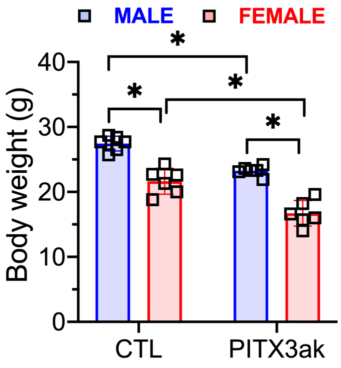


| | F (DFn, DFd) | |
|---|---|---|
| **interaction** | F (1, 20) = 0.3153 | P=0.5807 |
| **Pitx3 mutation** | F (1, 20) = 51.40 | P<0.0001 |
| **Sex** | F (1, 20) = 94.15 | P<0.0001 |

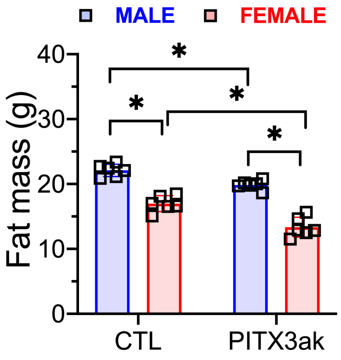


| F (DFn, DFd) | |
|---|---|
| F (1, 20) = 2.193 | P=0.1542 |
| F (1, 20) = 39.78 | P<0.0001 |
| F (1, 20) = 156.3 | P<0.0001 |

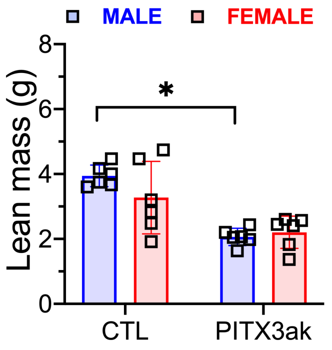


| F (DFn, DFd) | |
|---|---|
| F (1, 20) = 2.357 | P=0.1404 |
| F (1, 20) = 31.10 | P<0.0001 |
| F (1, 20) = 0.9988 | P=0.3295 |

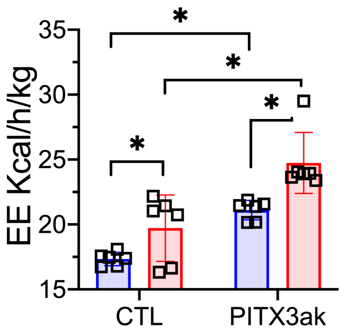


| | F (DFn, DFd) | |
|---|---|---|
| **interaction** | F (1, 20) = 0.7269 | P=0.4040 |
| **Pitx3 mutation** | F (1, 20) = 35.90 | P<0.0001 |
| **Sex** | F (1, 20) = 16.83 | P=0.0006 |

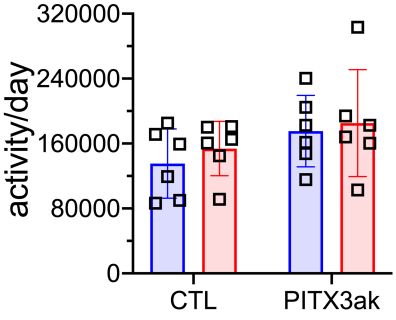


| F (DFn, DFd) | |
|---|---|
| F (1, 20) = 0.04951 | P=0.8262 |
| F (1, 20) = 3.320 | P=0.0834 |
| F (1, 20) = 0.5230 | P=0.4779 |

**Fig S2. Energy expenditure in male and female control (CTL) and Pitx3ak mice with food available ad-libitum in Experiment 1.** Waveforms show raw and smoother hourly data for each mouse, and group mean averages (top two panels).

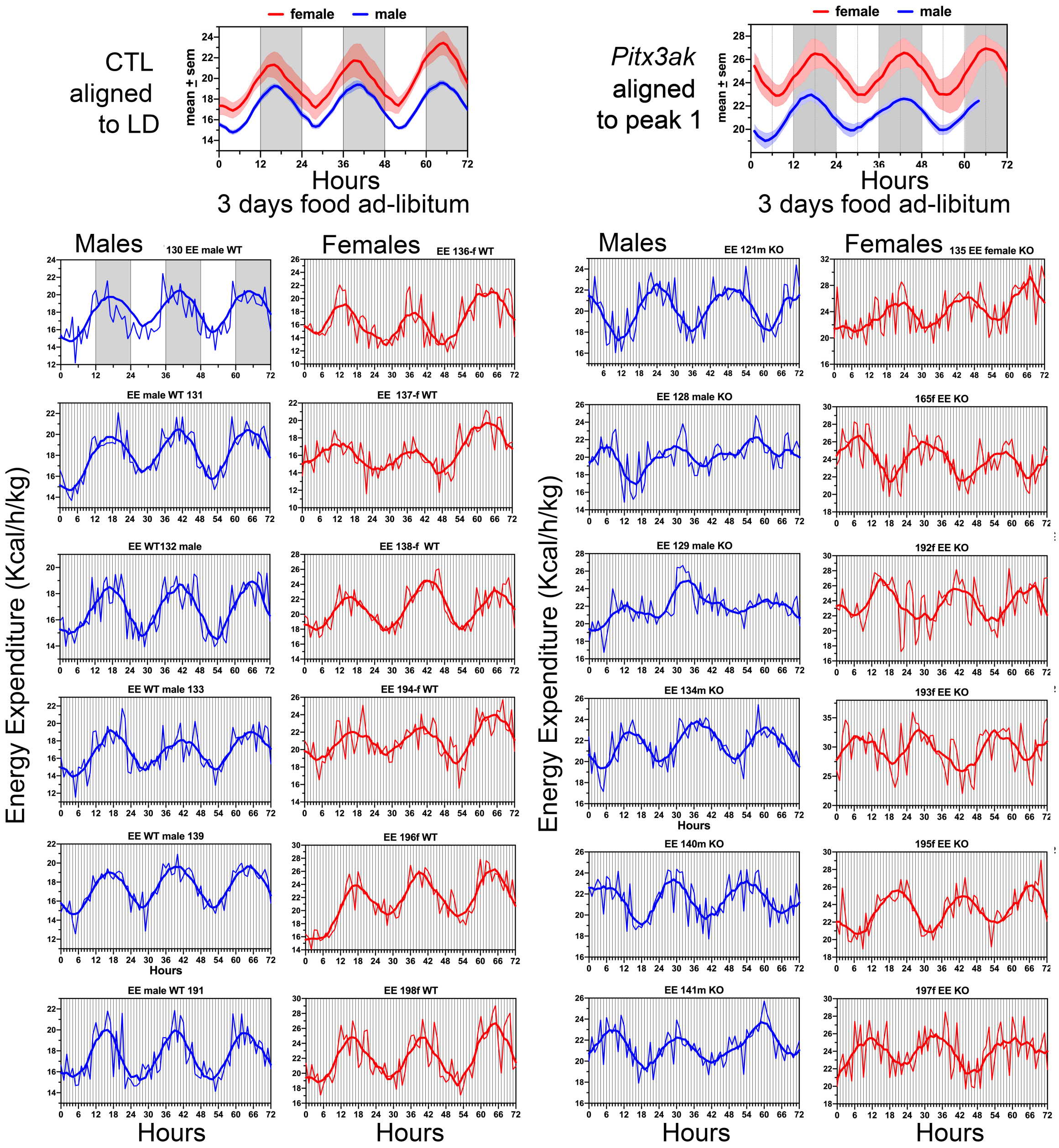

**Fig S3. Educed waveforms of locomotor activity for each *Pitx3ak* mouse with food available ad-libitum in Experiment 2.** The time series were folded modulo the free running period to create the waveform for each mouse (left column) which when averaged across mice eliminated the circadian periodicity (top left). Waveforms in the right column were aligned by the first peak of activity before averaging (top right).

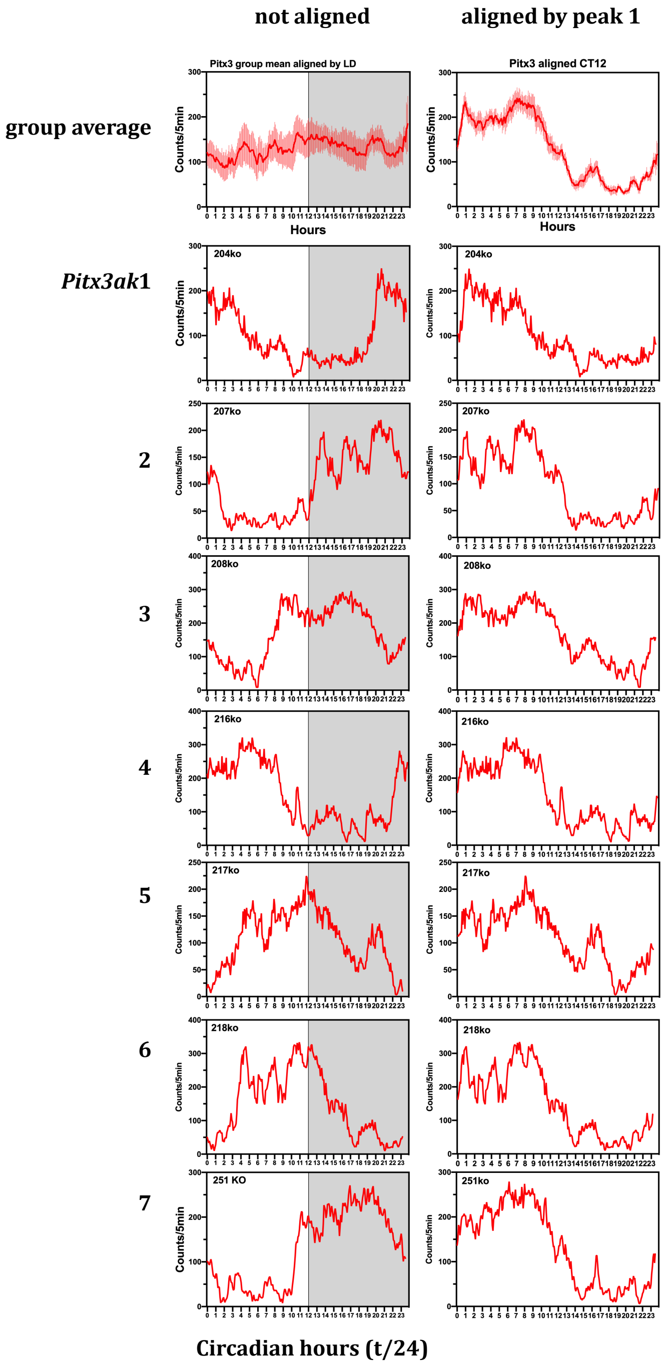
