## Supplemental Figures for "Mice hypomorphic for *Pitx3* define a minimal dopamine neuron population sufficient for entraining behavior and metabolism to scheduled feeding"

Fig S4. Survival of control (black) and Pitx3^ak^ (red) mice on 60% CR daily feeding. P value was computed using the log rank test.

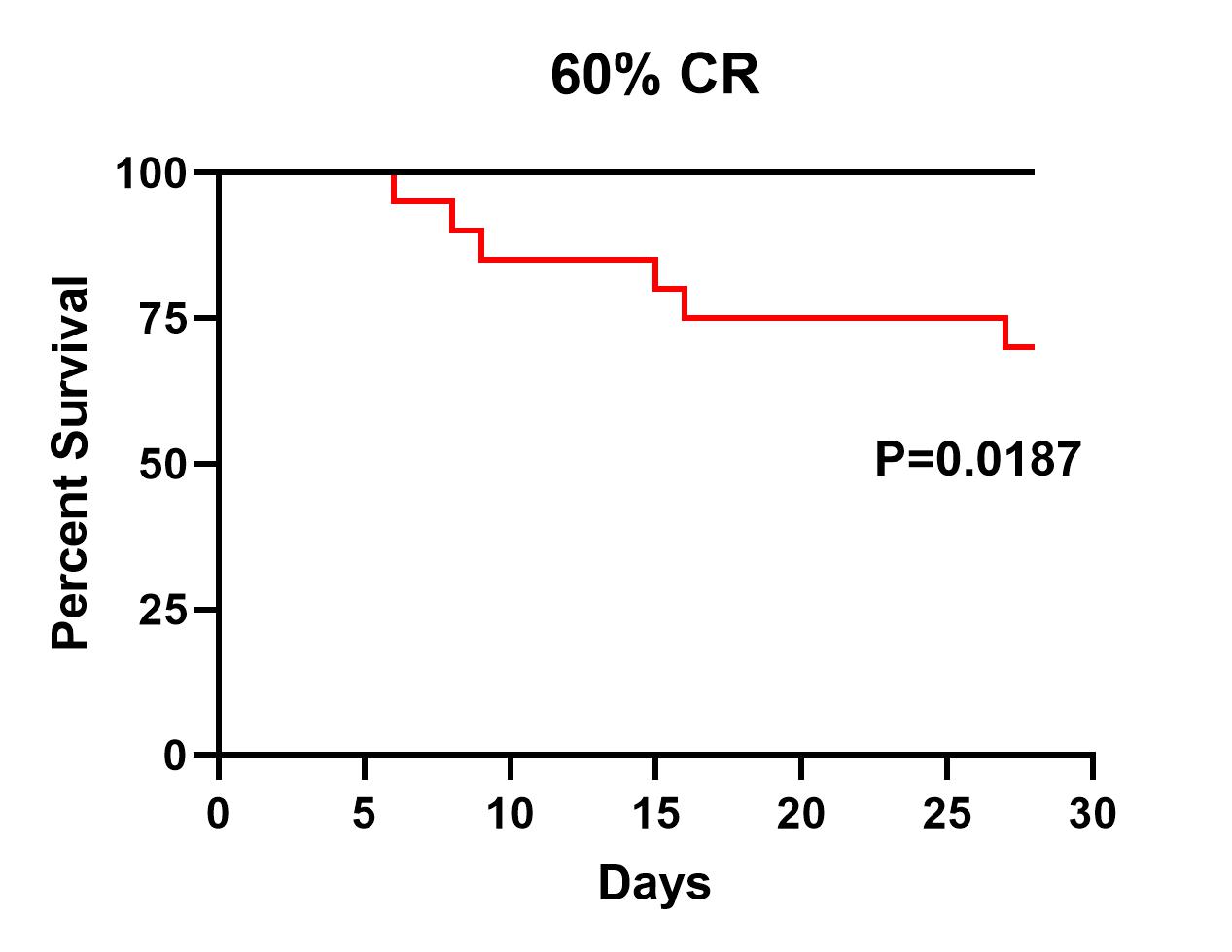
